# Generation of nonhuman primate retinitis pigmentosa model by *in situ* knockout of *RHO* in rhesus macaque retina

**DOI:** 10.1101/2020.07.29.226787

**Authors:** Shouzhen Li, Yingzhou Hu, Yunqin Li, Min Hu, Wenchao Wang, Yuqian Ma, Yuan Cai, Min Wei, Yichuan Yao, Yun Wang, Kai Dong, Yonghao Gu, Huan Zhao, Jin Bao, Zilong Qiu, Mei Zhang, Xintian Hu, Tian Xue

## Abstract

Retinitis pigmentosa (RP) is a form of inherited retinal degenerative disease that ultimately involves the macula, which is present in primates but not in the rodents. Therefore, creating nonhuman primate (NHP) models of RP is of critical importance to study its mechanism of pathogenesis and to evaluate potential therapeutic options in the future. Here we applied adeno-associated virus (AAV)-delivered CRISPR/SaCas9 technology to knockout the *RHO* gene in the retinae of the adult rhesus macaque (*Macaca mulatta*) to investigate the hypothesis whether non-germline mutation of the *RHO* gene is sufficient to recapitulate RP. Through a series of studies, we were able to demonstrate successful somatic editing of the *RHO* gene and reduced RHO protein expression. More importantly, the mutant macaque retinae displayed clinical RP phenotypes, including photoreceptor degeneration, retinal thinning, abnormal rod subcellular structures, and reduced photoresponse. Therefore, we suggest somatic editing of the *RHO* gene is able to phenocopy RP, and the reduced time span in generating NHP mutant accelerates RP research and expands the utility of NHP model for human disease study.

## Introduction

Retinitis pigmentosa (RP) is an inherited retinal disease, which initially shows signs of night blindness due to loss of rod photoreceptors, followed by loss of cone photoreceptors with reduced daytime vision [1, 2]. Over 50 genes are related to RP, including *RHO, PDE6B, RPE65*, and *CNGB1* (RETNET, https://sph.uth.edu/retnet/) [3-5]. To study the underlying mechanisms and develop therapeutic methods for RP, several RP mutant rodent models, including *rd1, rd10, Rho*^*-/-*^, and *Rho*^*P23H/+*^ mutant mice, have been established [6-9]. These models display progressive loss of photoreceptors and vision that mimic human RP disease [10]. However, rodent models are genetically and physically diverse from those of humans. In particular, rodent models lack macula and fovea in their retinae, which are important for high visual acuity in humans. In contrast, Nonhuman primates (NHPs) and humans share many genetic, anatomical, physiological, immunological, and behavioral similarities [11, 12]. Particularly, old world monkeys such as rhesus macaques have macula in retina. Considering that rodent models cannot fully recapitulate the pathogenesis of human RP and NHPs have a highly developed neuroanatomy closely resembling that of humans, an NHP RP model is highly desired.

Recently, NHP models of human diseases were generated by germline gene modification [13-17]. However, the generation of these NHP models is costly and technically challenging due to the difficulty of handling germline cells and monkey breeding. As such, the advancement of adeno-associated virus (AAV) delivered clustered regularly interspersed short palindromic repeats (CRISPR)/Cas9 for somatic gene editing appears to be a much more efficient and affordable way to generate NHP disease models. CRISPR/Cas9 requires a single-guide RNA (sgRNA) to guide Cas9 to generate double-stranded DNA breaks (DSBs) at target loci to introduce mutations [18]. Small insertions and deletions (indels) from nonhomologous end joining (NHEJ) DNA repair can disrupt the open reading frame (ORF) to knockout genes. It is worth noting that AAV-delivered CRISPR/Cas9-mediated gene editing can greatly benefit tissue- and region-specific studies by enabling selective manipulation of gene expression in specific tissue regions.

Rhodopsin, which is encoded by the *RHO* gene, is the first protein found to be mutated in RP [19, 20]. It is a G-protein coupled receptor (GPCR) located in the outer segment membranes of rod photoreceptors [21]. Furthermore, it is critical for phototransduction, converting light into electrical signals. *RHO* is the main causative gene for autosomal-dominant (adRP) and autosomal-recessive RP (arRP), accounting for 30%–40% of adRP cases and 8%–10% of all RP [22]. To model the most common adRP disease with loss of *RHO*, and to investigate whether non-germline mutation of the *RHO* gene is sufficient to recapitulate RP, we used AAV-delivered CRISPR/*Staphylococcus aureu*s Cas9 (SaCas9)-mediated gene editing to knock out the *RHO* gene in the rod photoreceptors of macaque (*Macaca mulatta*) *in vivo*. The *RHO*-knocked out retinae displayed typical RP photoreceptor degeneration, and mimicked human RP in regard to molecular, morphological, and physiological changes. Our study validated that AAV-delivered CRISPR/SaCas9 could somatically knockout a gene in a tissue-specific manner in adult macaques. This primate RP model could be a valuable tool to study the mechanisms and pathogenesis of human RP as well as to develop therapeutic strategies for the disease. In addition, the reduced time span in generating NHP mutant can accelerate RP research and expands the utility of NHP model for human disease research.

## Results

### *In vitro* knockout of *RHO* by CRISPR/SaCas9-mediated gene editing

To generate a *RHO*-knocked out macaque RP model based on CRISPR/Cas9 gene editing, three sgRNAs (sgRNA1, 2, 3) targeting the first exon of the *RHO* gene were designed to obtain a high rate of complete gene knockout (Fig. 1A). To construct Cas9 and sgRNA into one vector for high co-transduction rate and to overcome the package limitation of AAV (∼4.85 kb including both inverted terminal repeats (ITRs)), we chose SaCas9 for AAV delivery as it is ∼1 kb shorter than conventional *Streptococcus pyogenes* Cas9 (SpCas9). We generated the plasmid AAV-hSyn-SaCas9-U6-sgRNA using the human synapsin I (hSyn) promoter (448 bp) to drive neuron-specific expression of SaCas9 and make the plasmid 4.72 kb in length suitable for AAV package (Fig. 1B). Each sgRNA under the control of the U6 promoter was individually cloned with SaCas9. Plasmid AAV-hSyn-SaCas9 containing SaCas9 and a scramble sgRNA was used as the control. To make sure that hSyn promoter can work efficiently in both HEK293T cells *in vitro* and photoreceptors *in vivo*, we transfected HEK293T cells with AAV-hSyn-mKate2 (Fig. S1A), and produced AAV virus to infect mouse photoreceptors by subretinal injection *in vivo* (Fig. S1B), respectively. The results indicated that hSyn promoter could efficiently drive the gene expression in the two systems.

**Figure 1:**
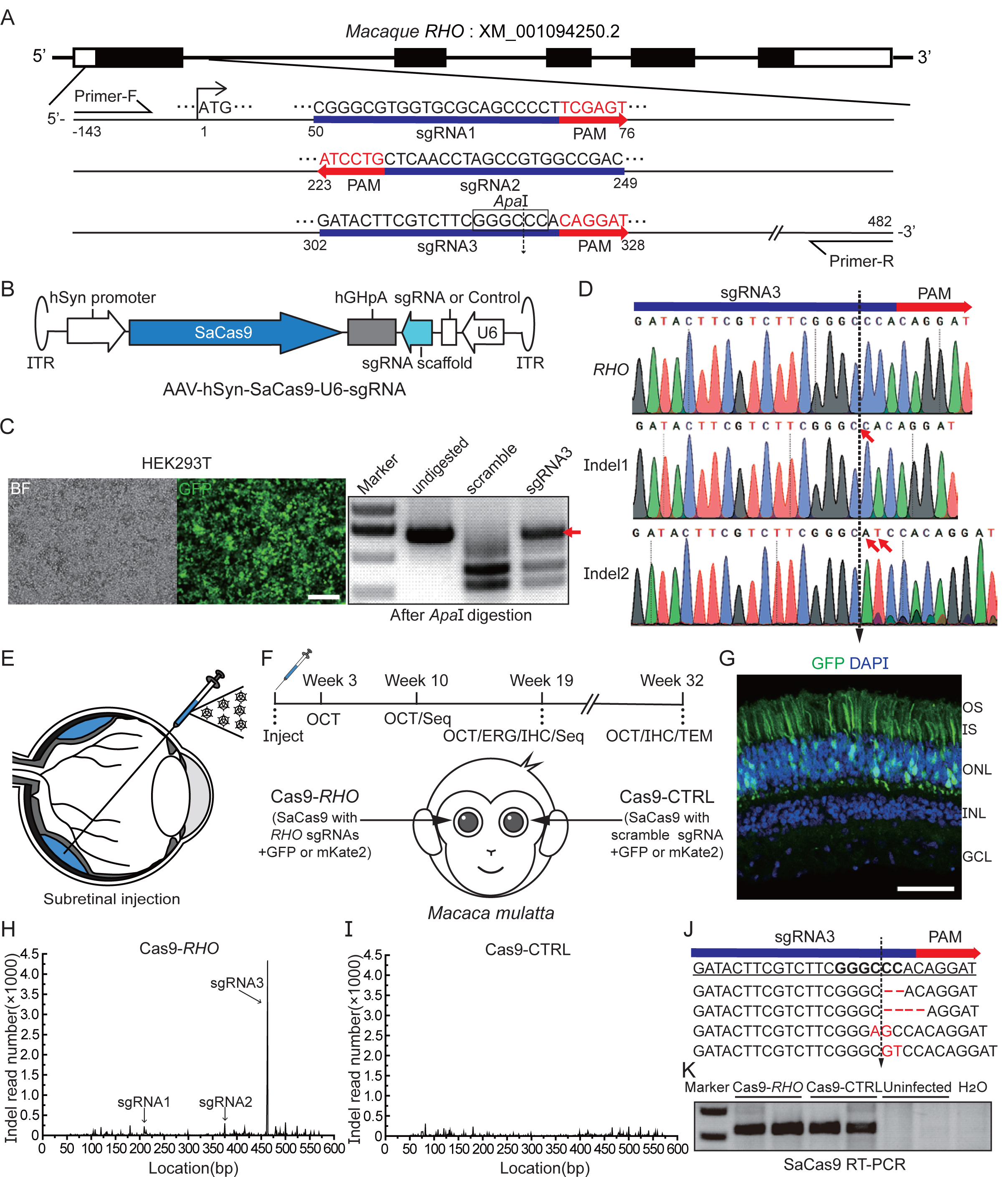
Knock-out of *RHO* gene in rhesus macaques via CRISPR/SaCas9 *in vitro* and *in vivo*. (A) Design of sgRNAs at the first exon of macaque *RHO* gene. Three sgRNAs were designed to target the first exon of *RHO*, with 52 bp and 146 bp intervals based on PAM sequence of SaCas9. Sequence of each sgRNA (blue bar) is shown with subsequent PAM sequence (red bar). *Apa*I digestion site exists in predicted cleavage site of SaCas9-sgRNA3. Two primers spanning first exon of *RHO* were designed for PCR amplification. (B) Schematic representation of AAV-hSyn-SaCas9-U6-sgRNA vector. Human synapsin I promoter drives the expression of SaCas9 and U6 promoter drives the expression of sgRNA in the same vector. (C) Transfection efficiency of AAV-CBh-GFP vector in HEK293T cells (left panel). Scale bar, 80 μm. After *Apa*I digestion, PCR products of the first exon were analyzed. A band at 670 bp indicated the occurrence of indels in sgRNA3 targeting site (right panel, red arrow). (D) Sequencing of 670 bp band indigestible by *Apa*I showed the existence of indels in *RHO* genome: i.e., 1 bp deletion occurred in Indel 1 sample and 2 bp insertion occurred in Indel 2 sample. (E) Schematic representation of subretinal injection in macaques. (F) Diagram of entire experimental design. A series of analyses were carried out over 32 weeks. (G) Viral transduction of AAV-CBh-GFP in macaque retinae. Scale bar, 50 μm. (H) Number of indel-existing reads (×1000) at each base pair located in first exon of *RHO* in Cas9-*RHO* retinae. There was an obvious peak at sgRNA3 predicted targeting site. (I) Number of indel-existing reads (×1000) at each base pair location in Cas9-CTRL retinae. (J) Representative sequences of indel-existing reads at sgRNA3 targeting site. (K) RT-PCR results of SaCas9 expression in Cas9-*RHO*, Cas9-CTRL, uninfected retinae, and distilled water (H2O). OS, outer segment; IS, inner segment; ONL, outer nuclear layer; INL, inner nuclear layer; GCL, ganglion cell layer.

To examine the efficiency of the *RHO* gene editing by the CRISPR/SaCas9 system *in vitro*, sgRNA3, which is homologous to the human *RHO* sequence, was tested in human HEK293T cells. Plasmid AAV-hSyn-SaCas9-U6-sgRNA3 was transfected into cells (Cas9-*RHO* hereafter), together with an AAV-CBh-GFP vector (Fig. S1C) for evaluating transfection efficiency (Fig. 1C, left panel). Plasmids AAV-hSyn-SaCas9 and AAV-CBh-GFP were co-transfected as the control (Cas9-CTRL hereafter). As the SaCas9/sgRNA3 predicted cutting site overlapped with an *Apa*I restriction enzyme site, the *RHO* sequence with indels could escape from the digestion of *Apa*I (Fig. 1A, and C right panel). Primers spanning the first exon of *RHO* were designed to amplify the corresponding DNA sequence and the polymerase chain reaction (PCR) products were digested by *Apa*I. There was a clear DNA band that could not be cut by *Apa*I in the Cas9-*RHO* group, suggesting the occurrence of indels in *RHO* (Fig.1C, right panel). The cleavage efficiency of sgRNA3 was ∼50% *in vitro*, which was estimated by dividing the uncleaved band intensity to the total DNA band intensity, followed by correction using the HEK293T transfection rate using GFP signals. The indels were further identified by sequencing of the *Apa*I undigested band, which lead to ORF shifts in *RHO* sequencing (Fig. 1D). Thus, CRISPR/SaCas9 could efficiently knockout *RHO* gene *in vitro*.

### *In vivo* knockout of *RHO* by AAV-delivered CRISPR/SaCas9 system in rhesus macaque retinae

ShH10 is an AAV serotype that has been developed to target müller glial cells in the retina by intravitreal injection [23]. Interestingly, serotype AAV/ShH10 can also effectively infect mouse photoreceptor by subretinal injection [24]. We further confirmed that AAV/ShH10-hSyn-mKate2 effectively infected the mouse and macaque photoreceptors after subretinal injection (Fig. S1B and D). Thus, we packaged all AAV vectors using serotype ShH10 for macaque retinal transduction. The eyes of macaques were through subretinal injection with a modified Hamilton needle (38 G, 40 G) passing through the vitreous body (Fig. 1E). Three injection sites were chosen for each eye, and 50 μl of AAV was injected at each injection site. The right retinae infected with the combination of AAV/ShH10-hSyn-SaCas9-U6-sgRNA1, 2, 3 were designated as the Cas9-*RHO* group, and the left retinae infected with AAV/ShH10-hSyn-SaCas9 were designated as the Cas9-CTRL group (Fig. 1F). Either AAV/ShH10-CBh-GFP or AAV/ShH10-CBh-mKate2 was co-injected to indicate the virus-infected area and its transduction efficiency (Fig. 1G). About 10-20% of the retina was infected by AAV. In four retinae, mKate2 was used instead of GFP as GFP can interfere with sodium fluorescein signals in the fundus fluorescence angiography (FFA) (Fig. S1C). A series of analyses was performed within 32 weeks after injection to examine the phenotypes of infected retinae, including optical coherence tomography (OCT), FFA, *ex vivo* electroretinography (ERG), and transmission electron microscopy (TEM) (Fig. 1F). Altogether, four rhesus macaques were analyzed for this study (Table 1).

**Table 1.** Summary of *RHO* KO in rhesus macaques after subretinal AAV administration.

| Animal No. | Gender | Age at dosing<br>(years) | Retina | AAV | Titer<br>(vg/ml) | Volume | Sacrificed time<br>Post injection | Experiments<sup>a</sup> |
| --- | --- | --- | --- | --- | --- | --- | --- | --- |
| 1 | M | 13.2 | Left | PBS | No | 50μl×3 | Week 26 | ERG, IHC, TEM |
|  |  |  | Right | Cas9-*RHO* + GFP | 5×10<sup>12</sup> | 50μl×3 |  |  |
| 2 | M | 12.2 | Left | Cas9-CTRL + GFP | 5×10<sup>12</sup> | 50μl×3 | Week 19 | ERG, Seq, IHC, WB |
|  |  |  | Right | Cas9-*RHO* + GFP | 5×10<sup>12</sup> | 50μl×3 |  |  |
| 3 | M | 8.2 | Left | Cas9-CTRL + mKate2 | 5×10<sup>12</sup> | 50μl×3 | Week 32 | OCT, FA, IHC, TEM, WB |
|  |  |  | Right | Cas9-*RHO* + mKate2 | 5×10<sup>12</sup> | 50μl×3 |  |  |
| 4 | M | 8.2 | Left | Cas9-CTRL + mKate2 | 5×10<sup>12</sup> | 50μl×3 | Week 10 | Seq, RT-PCR |
|  |  |  | Right | Cas9-*RHO* + mKate2 | 5×10<sup>12</sup> | 50μl×3 |  |  |

<sup>a</sup>ERG, electroretinogram; IHC, immunohistochemistry; TEM, transmission electron microscope; Seq, PCR deep sequence; WB, western blot; OCT, optical coherence tomography; FA, fluorescein angiography; RT-PCR, SaCas9 reverse transcript PCR.

To examine whether *RHO* was knocked out *in vivo*, we collected GFP-positive region in retina and extracted the genomic DNA for further molecular analysis. The PCR products amplified from the first exon of *RHO* were used for *de novo* sequencing to examine whether *RHO* was successfully knocked out (Fig. 1H-J). The number of indel-existing reads was quantified in both the Cas9-*RHO* and Cas9-CTRL groups. Results revealed significant indel-existing reads at the desired location, between the 18^th^ and 19^th^ base pair of sgRNA3 (Fig. 1H and J). These indels could result in an ORF shift and may produce dysfunctional RHO proteins. In contrast, sgRNA1, 2 showed less effective gene editing with fewer indels compared with that of sgRNA3 (Fig.1H). The expression of SaCas9 was investigated in virus-infected areas by reverse transcription-PCR (RT-PCR) (Fig. 1K). The detection of SaCas9 suggested the successful delivery and expression of SaCas9 in the rhesus macaque retinae. To examine the potential off-target loci from the CRISPR/Cas9 technique, we sequenced the PCR product at sgRNAs potentially targeted regions. We identified no genetic mutations in these potential target sites, as predicted by online tools (http://www.rgenome.net/cas-offinder/) (Fig. S2 and Table 2). Thus, the AAV-delivered CRISPR/SaCas9 system successfully edited the *RHO* gene in adult rhesus macaque retinae without off targeting effects.

**Table 2.**
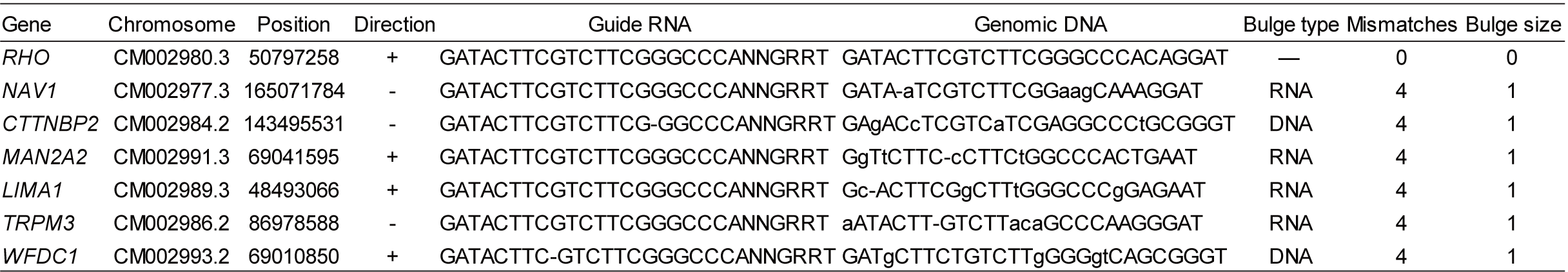
Summary of potential off-target sites from macaque coding sequence predicted by cas-offinder software.

### Knockout of *RHO* by AAV-CRISPR/SaCas9 generated RP-like photoreceptor degeneration

To investigate the morphological changes in rod photoreceptors from Cas9*-RHO* retinae, the rod outer segments were visualized by rhodopsin staining at post injection week (PIW) 19 (Fig. 2A-C). The length from the photoreceptor cell nuclei to the RHO-positive end, including both inner and outer segments, was calculated in both Cas9-CTRL and Cas9-*RHO* retinae. Compared with uninfected areas, the photoreceptor inner/outer segments of Cas9-CTRL-infected regions showed no significant differences in morphology (Fig. 2B). In the virus-infected Cas9-*RHO* retinae, however, the photoreceptor inner/outer segments displayed distinct degrees of degeneration (Fig. 2C). In some regions of the Cas9-*RHO* retinae, almost no rhodopsin expression was detected (Fig. 2C, right). The length of the photoreceptor inner/outer segments in the Cas9-*RHO* retinae was reduced to ∼47% of that in the Cas9-CTRL retinae (Fig. 2D). We then studied whether the RHO protein level was changed by western blot analysis (Fig. 2E). Results indicated that the expression level of RHO in Cas9-*RHO* retinae was reduced to ∼45% of that in the Cas9-CTRL retinae (Fig. 2F), thus suggesting a loss of RHO in the mutant retinae.

**Figure 2:**
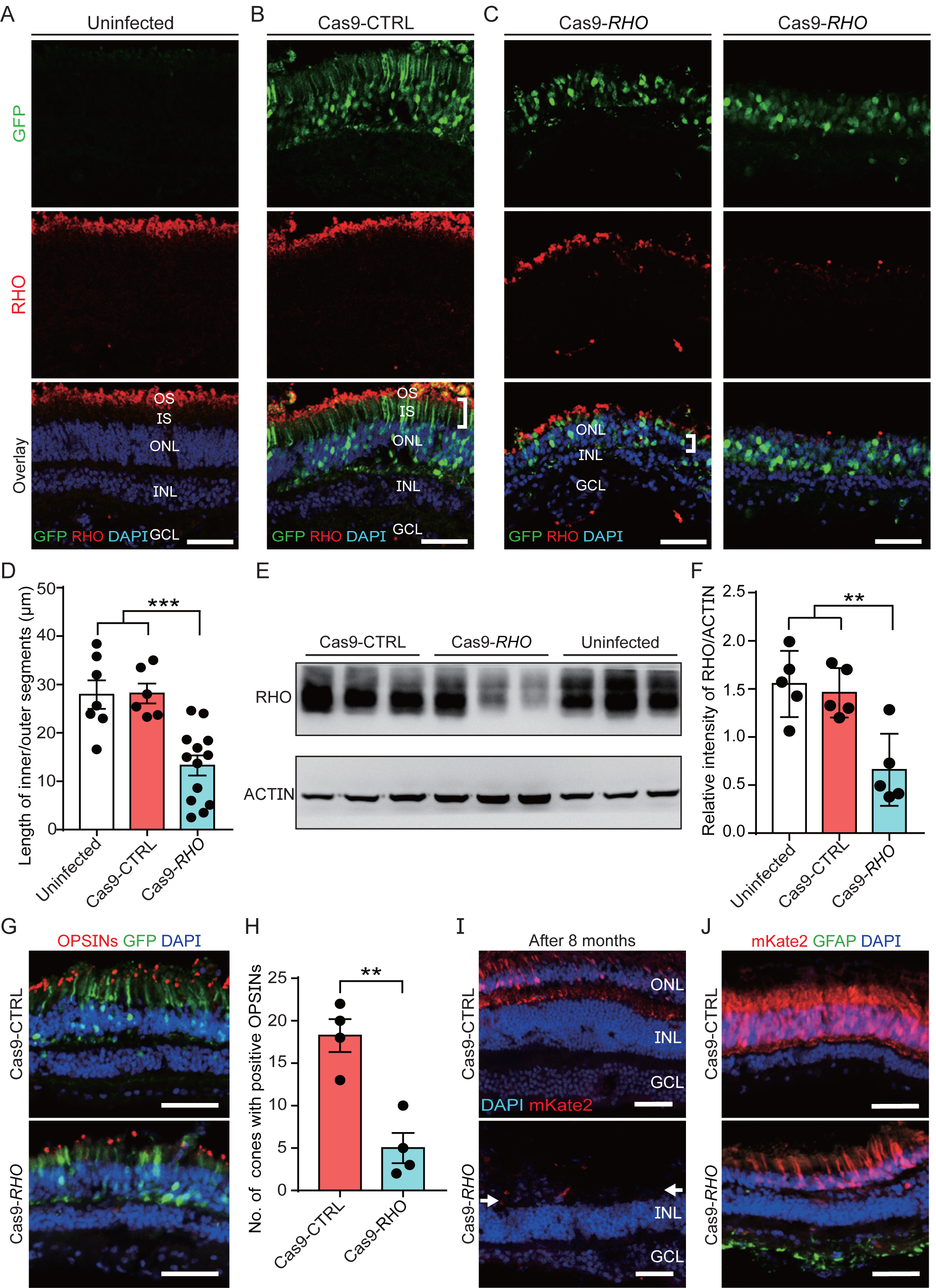
Morphology of photoreceptors in Cas9-*RHO* macaque retinae. (A) Immunostaining of RHO in retinae without AAV transduction. RHO, rhodopsin. (B) Immunostaining of RHO in Cas9-CTRL retinae. Cas9-CTRL retinae showed normal morphology, with regular inner segments and rhodopsin expression. GFP, AAV-CBh-GFP as a reporter. (C) Immunostaining of RHO in Cas9-*RHO* retinae. Photoreceptors in infected area showed shorter inner/outer segments and reduced rhodopsin expression. In some areas of Cas9-*RHO* retinae, the expression of RHO and the inner/outer segments of photoreceptors were all lost (right panel). (D) Quantification of inner/outer segment length in uninfected region of retinae, Cas9-CTRL and Cas9-*RHO* retinae. n = 7–13 slices from two macaques for uninfected and Cas9-*RHO* retinae, n = 6 slices from one macaque for Cas9-CTRL retinae. \*\*\**P* < 0.001, one-way ANOVA. (E) Western blot showed less RHO expression in Cas9-*RHO* retinae compared with Cas9-CTRL or uninfected retinae. ACTIN was used as internal control. (F) Quantification of western blots showed RHO expression significantly decreased in Cas9-*RHO* retinae compared with Cas9-CTRL and uninfected retinae. n = 5 samples from two macaques of either Cas9-CTRL or Cas9-*RHO*. \*\**P* < 0.01, one-way ANOVA. (G) Cone staining with OPN1-LW/MW and OPN1-SW in Cas9-CTRL and Cas9-*RHO* retinae. OPN1-LW/MW and OPN1-SW were co-stained. (H) Quantification of cones with positive OPSINs in per 200μm Cas9-CTRL and Cas9-RHO retinae. n = 4 slices from one macaque. **P < 0.01, unpaired Student’s t-test. (I) Cas9-CTRL retinae showed normal retinal layers after eight months. ONL nearly disappeared in Cas9-*RHO* retinae (white arrow), whereas INL and GCL showed normal thickness. mKate2, AAV-CBh-mKate2 as reporter. (J) Activated glia in Cas9-*RHO* retinae, labelled by GFAP, but very few GFAP signals in Cas9-CTRL retinae was detected. Scale bar, 50 μm. OS, outer segment; IS, inner segment; ONL, outer nuclear layer; INL, inner nuclear layer; GCL, ganglion cell layer.

Cone photoreceptors, which are important for daytime vision, were identified by the mixture of OPSIN antibodies staining against both long/mid-wave OPSIN (OPN1-LW/MW) and short-wave OPSIN (OPN1-SW) [25] (Fig. 2G). The number of cones with positive OPSINs immuno-activity in Cas9-*RHO*-infected areas was greatly reduced to ∼27% of that in the Cas9-CTRL-infected areas (Fig. 2H), suggesting cone photoreceptor degeneration in Cas9*-RHO* retinae. As RHO is specifically expressed in rod photoreceptors, the loss of cones with OPSINs in Cas9-*RHO* was possibly a secondary effect of the loss of rod photoreceptors [26]. It is worth mentioning that the outer nuclear layer (ONL) was completely lost in the macular region after eight months (Fig. 2I), indicating progressive photoreceptor degeneration. These results are consistent with age-dependent progressive retinal degeneration found in *Rho*^P23H/+^ rodent models, which show loss of most rods and cones in ventral retina at around three months [6]. Macroglial cells (astrocytes and Müller cells) are selectively activated in human RP retinae [27]. In Cas9-*RHO* retinae, the expression of glial fibrillary acidic protein (GFAP), a marker for reactive gliosis in the retina [28], was enhanced (Fig. 2J). Collectively, these findings demonstrated that Cas9-*RHO* retinae displayed typical photoreceptor degeneration with retinal gliosis.

### Cas9-*RHO* retinae showed clinical signs consistent with RP

To further analyze the retinopathy *in vivo*, we examined the Cas9-*RHO* retinae by FFA and OCT. All macaque retinae were confirmed to be morphologically normal by fundus photography and FFA before virus-injection (Fig. 3A). There were three retinal injection sites, covering the macula, superior temporal venule, and inferior temporal venule of the retina, respectively (Fig. 3A, right). OCT analysis after injection revealed that the retinal detachments caused by virus-injection recovered within three weeks (Fig. 3B), suggesting no significant long-term damage from the injections alone. FFA analysis showed that the Cas9-*RHO* retinae displayed hyperfluorescent flecks in the virus-infected areas, indicating the occurrence of retinal telangiectasia leakage (Fig. 3D). In the Cas9-CTRL retinae, despite a few localized glial scars in the retinae caused by needle damage, very few hyperfluorescent flecks were detected at the infected region (Fig. 3C).

**Figure 3:**
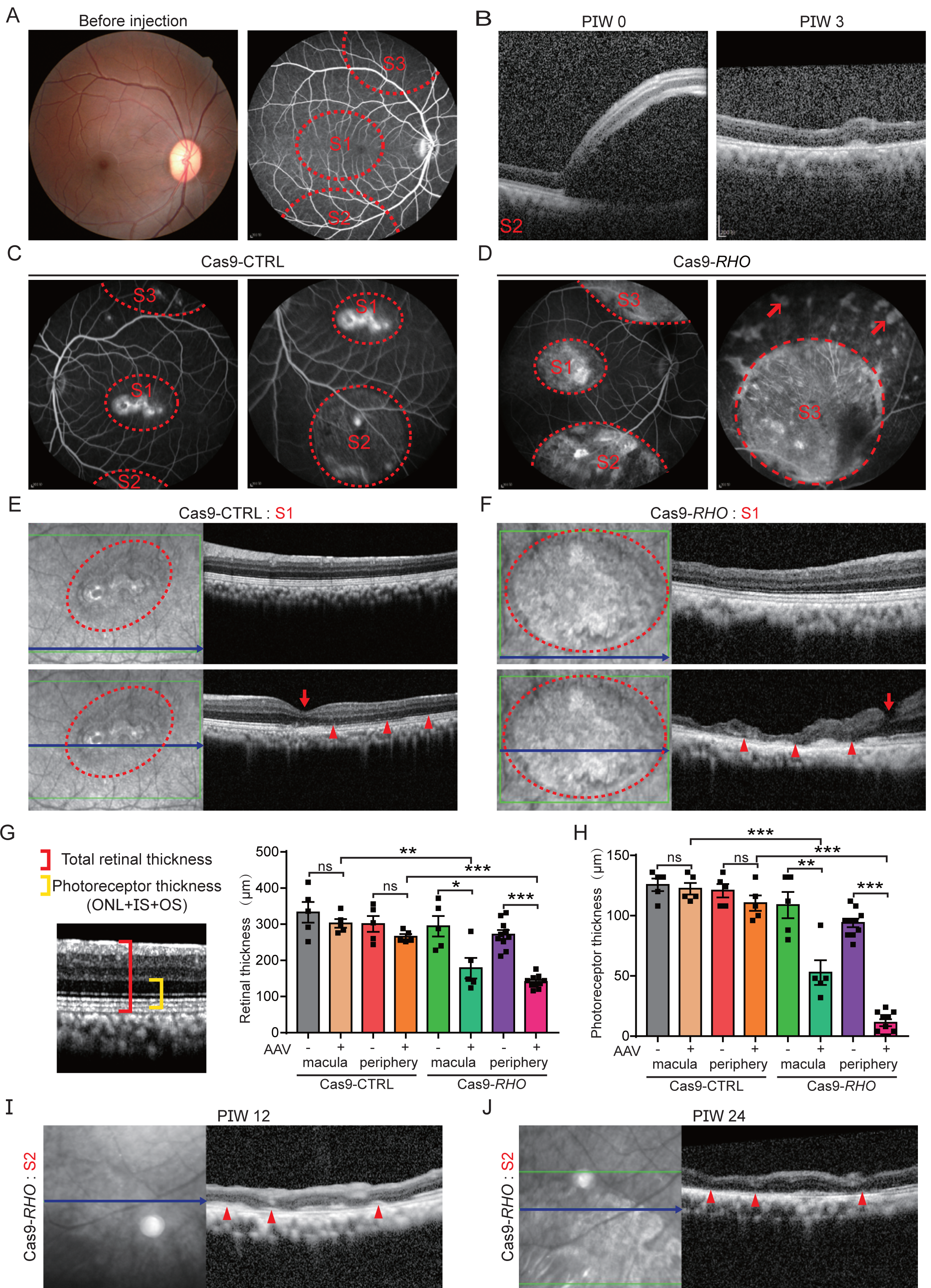
Pathological changes in Cas9-*RHO* macaque retinae. (A) Fundus images (left) and FFA (right) showed normal retinal morphology of macaques before injection. Diagram of subretinal injection sites highlighted by red ring (right). S, injection site. (B) AAV injection bulge (left) was absorbed (right) within PIW 3. (C) FFA images of Cas9-CTRL retinae showed a few localized glial scars at injection site. (D) FFA images of Cas9-*RHO* retinae showed dramatic hyperfluorescent flecks in infected areas (red arrow). (E) OCT images of uninfected (above) and infected areas (below) at S1 in Cas9-CTRL retinae showed normal retinal structure. Blue line indicates scanned site and red ring indicates infected area. Red arrow indicates fovea centralis. Red triangle indicates photoreceptor segments (ellipsoid zone or interdigitation zone). (F) OCT images of S1 in Cas9-*RHO* retinae showed structure of ONL was severely degenerated in infected area, but normal in nearby uninfected area. (G) The diagram of statistics of total retinal thickness (red) and photoreceptor thickness (yellow) in OCT images (left). The total retinal thickness of Cas9-*RHO* and Cas9-CTRL retinae was quantified (right). +, the AAV infected area; -, the AAV-uninfected area; \**P*<0.05, \*\**P <* 0.01, \*\*\**P <* 0.001, unpaired Student’s t-test. (H) Quantification of the thickness of photoreceptors of Cas9-*RHO* and Cas9-CTRL retinae. (I) and (J) Progressive loss of photoreceptor segments (red arrow) in Cas9-*RHO* retinae from PIW 12 to PIW 24.

To obtain noninvasive cross-sectional retinal images for the diagnosis of RP, we performed OCT to evaluate the total retinal thickness and photoreceptor thickness of the macaque retinae at 6 months post AAV administration (Fig. 3E-H, Fig.S3A-B). In the Cas9-CTRL retinae, the infected ONL exhibited no significant changes in total retinal thickness and photoreceptor thickness in either the macula or the peripheral regions compared with nearby uninfected regions (Fig. 3E, G and H). The presence of an ellipsoid zone is indicative of normal functioning photoreceptors (Fig. 3E and Fig.S3A). In the Cas9*-RHO* retinae, however, the ellipsoid zone was either disrupted or even absent (Fig. 3F and Fig.S3B). The total retinal thickness and photoreceptor thickness of the infected macula and periphery Cas9*-RHO* retinae were significantly decreased (Fig. 3F, G and H), suggesting potential vision loss in mutant macaque retinae. Moreover, the mutant retinae showed progressive degeneration as the affected retinae became increasingly thinner from PIW 12 to 24 (Fig. 3I and J). These results validate the RP retinal thinning observed in clinical diagnosis. Collectively, the *RHO*-mutated macaque retinae developed RP pathological lesions, thereby phenocopying human RP disease.

### Abnormal subcellular structure of photoreceptors in Cas9-*RHO* retinae by TEM analysis

The subcellular structures of photoreceptors were further evaluated by TEM analysis. The Cas9-CTRL retinae displayed nicely arranged rods with normal subcellular morphology (Fig. 4A), i.e., long dense mitochondria in the inner segments (Fig. 4B), regular intact discs in the outer segments (Fig. 4C), and regular nuclei with a high nuclear-cytoplasmic ratio (Fig. 4D). In the Cas9-*RHO* retinae, however, we observed vacuolated, even fragmented, mitochondria in the inner segments (Fig. 4E and F) and some disrupted and shortened outer segments scattered in the ONL with massive disorganization of membrane discs (Fig. 4E and G). Vacuolated mitochondria have been observed in human RP photoreceptors, suggesting that this model mimicked the subcellular pathological changes in human RP [29]. Previous studies in mice suggest that *RHO* is involved in rod disc formation [30]. Here, the Cas9-*RHO* rhesus macaque retinae demonstrated shortened and disorganized rod discs. Moreover, the photoreceptors in Cas9-*RHO* retinae exhibited strong cell apoptosis (Fig. 4H). Taken together, the *RHO*-mutant macaque photoreceptors displayed typical RP defective subcellular structure and cellular apoptosis.

**Figure 4:**
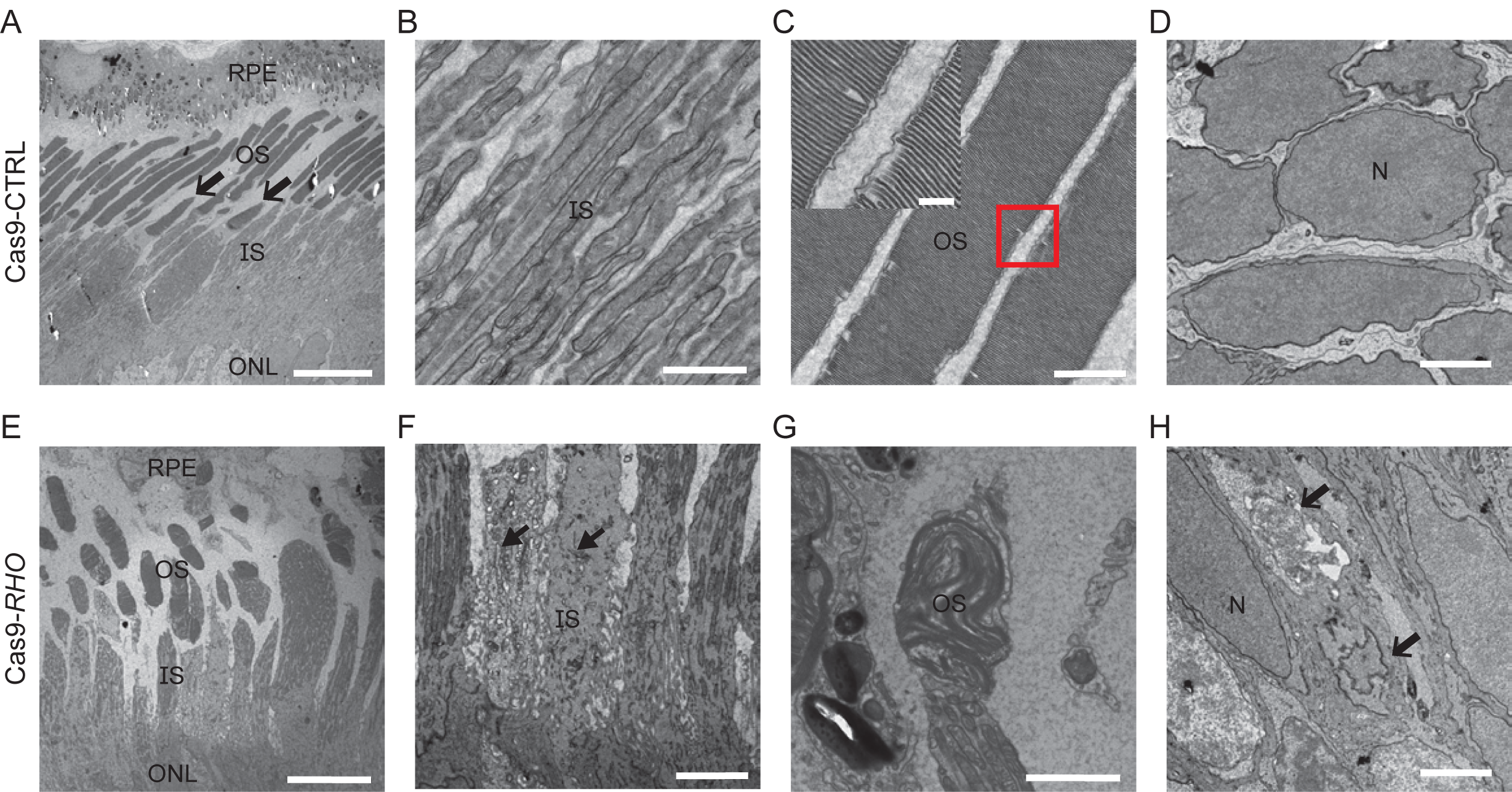
TEM analysis of photoreceptors in Cas9-*RHO* and Cas9-CTRL retinae. (A) TEM image of the photoreceptors in Cas9-CTRL retinae. Black arrow, cone photoreceptors. (B) Zoomed-in image shows the dense and long mitochondria in inner segments in Cas9-CTRL retinae. (C) Details of the outer segments in Cas9-CTRL retinae, showing entire membrane discs. (D) TEM image of Cas9-CTRL retinae shows normal morphology of photoreceptor soma in ONL, which displayed high nuclear-cytoplasmic ratio. (E) TEM image of the photoreceptors in Cas9-*RHO* retinae. (F) Zoomed-in image shows the vacuolated and degenerated mitochondria in the inner segments (black arrow). (G) Disrupted and shortened outer segments existed in Cas9-*RHO* retinae. (H) TEM image of Cas9-*RHO* retinae shows disrupted and atrophic cell nuclei in outer nuclear layer (open arrow). RPE, retinal pigment epithelium; OS, outer segment; IS, inner segment; ONL, outer nuclear layer; N, nucleus. Scale bar, 10 μm in A and E; 1 μm in B, C, D, and H; 2 μm in F and G.

### Impaired retinal physiological function in Cas9-*RHO*-mutant retinae

To determine the effect of *RHO* mutation on retinal physiological functions, *ex vivo* ERG recordings were performed in both Cas9-CTRL and Cas9-*RHO* retinae. Due to the virus-infected area covering only a small portion (∼10-20%) of the whole retina, *in vivo* ERG may not detect changes between Cas9-CTRL and Cas9-*RHO* retinae. Thus, *ex vivo* ERG was chosen to specifically record the infected area (Fig. 5A) [2]. As previously reported, *ex vivo* ERG in mouse (Fig. 5B) displays different waveforms compared with *in vivo* ERG and rhesus macaque retinae (Fig. 5D and E), with prolonged a-waves intermingled with b-waves [31]. A series of 20-ms light flashes from 3.8 to 1 515.6 photons•μm^-2^ was used to stimulate photoreceptors. In the Cas9-CTRL retinae, no significant differences in photoresponse were observed between the infected and adjacent uninfected areas (Fig. 5C-E). In contrast, the infected areas in the Cas9-*RHO* retinae showed significantly decreased photoresponse (Fig. 5F-H). In some severely degenerated areas, no ERG signals could be recorded. Therefore, loss of *RHO* impaired retinal electrophysiological function in rhesus macaque retinae.

**Figure 5:**
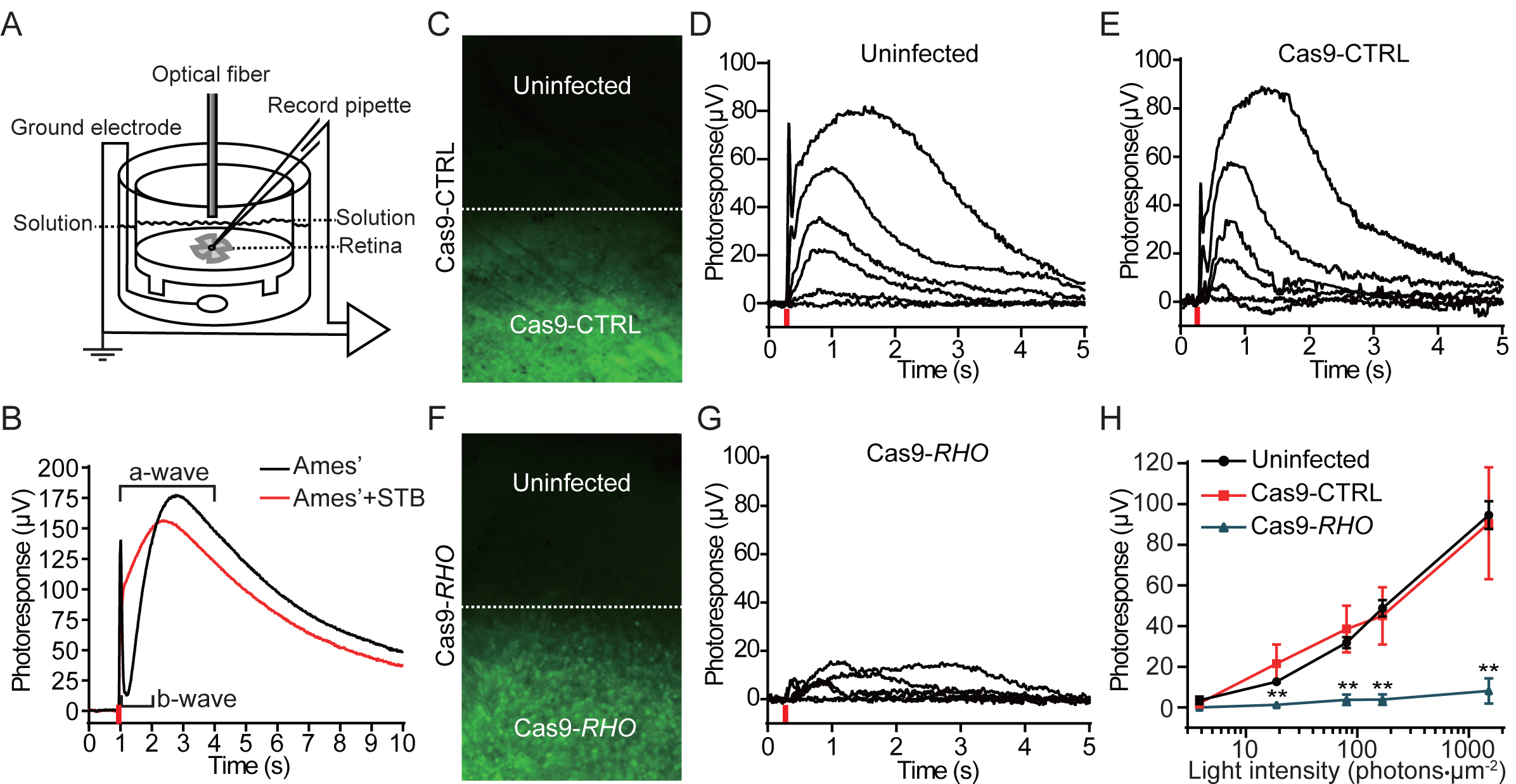
*Ex vivo* ERG recordings of uninfected, Cas9-*RHO*, and Cas9-CTRL retinae at four months post AAV injection. (A) Schematic representation of *ex vivo* ERG. (B) ERG recording in mice under Ames’ solution with or without synaptic transmission blockers (STBs). Prolonged a-wave was intermingled with b-wave. Intensity of light stimulation (red line) was 8 × 10^4^ photons/μm^2^. (C) Diagram of ERG recording regions in Cas9-CTRL retinae. (D) Traces of photoresponse under gradient light intensities in uninfected retina. Red line shows time of light stimuli. (E) Traces of photoresponse under gradient light intensities in Cas9-CTRL retinae. (F) Diagram of ERG recording in Cas9-*RHO* retinae. (G) Traces of photoresponse under gradient light intensities in Cas9-*RHO* retinae. (H) Quantification of ERG amplitudes in uninfected, Cas9-CTRL, and Cas9-*RHO* retinae under gradient light intensities. n = 20 retinal pieces from two macaques in uninfected group; n = 2 retinal pieces from one macaque in Cas9-CTRL group; n = 7 retinal pieces from two macaques in Cas9-*RHO* group. \*\**P* < 0.01, unpaired student’s *t*-test.

## Discussion

In this study, we generated a primate RP model with somatic knockout of *RHO* using the AAV-CRISPR/SaCas9 system. By using a series of molecular, cellular, and physiological analyses, we suggest the Cas9-*RHO* retinae phenocopied RP disease, which showed loss of RHO protein, photoreceptor degeneration, thinning retinae, and reduced physiological functions. We therefore successfully established a primate RP model for studying human RP pathology and developing therapeutic strategies.

Rodent RP models were widely used in RP mechanistic study, however, rodent models are genetically and physically far from humans, and lack of macula and fovea. Thus, the macaque RP model is more appropriate for studying its mechanism of pathogenesis and evaluating potential therapeutic options in the future. Compared with previous swine and canine RP models [32, 33], non-human primate we used here is *Macaca mulatta* evolutionarily closer to humans. Thus, our model may show phenotypes that are more similar to human RP in macular degeneration. Very recently, there have been reports of a primate with a very rare form of achromatopsia [34] and a report of Bardet-Biedl Syndrome in macaques [35]. However, those primate models mimic rare types of retinal degeneration. In contrast, *RHO* mutant is one of the most prevalent genes in the human RP, thus, the Cas9-*RHO* is more representative for retinal degeneration in human RP disease.

RP is a chronic, progressive, hereditary, dystrophic retinal degeneration. In this way, using germline transmission to establish an etiological RP could be able to better reflect the occurrence and development of the disease, and provide a more reliable model for future treatment. However, due to the technical difficulties, only a few labs in the world can achieve germline transmission in generating NHP models [13, 17, 36, 37]. Thus, it is not realistic to encourage the wide usage of germ cell gene editing in monkeys for modeling human diseases over the scientific community. In contrast, AAV-mediated CRISPR/SaCas9 is a rapid and affordable approach to generate NHP models for large-scale analysis of human RP phenotypes and mechanisms. Moreover, mutant retinae can be generated alongside control retinae, which provides a precise internal control for phenotypic analysis of mutant retinae. In our RP model, the virus-infected area is about 10-20% in whole retina. To increase the gene-editing efficiency, the injection of AAV-virus at more injection sites and more time points are recommended for the future study. In addition, more sgRNAs can be tested, and cloned all together in one vector to achieve high efficiency in the future.

*RHO*-adRP patient phenotype can be grouped into two major categories [38, 39]. Patients in class A have severe loss of rods from early life, and patients in class B can have rods that survive for decades into late adult life in some retinal regions. In our study, the morphological and physiological features of the Cas9-*RHO* model closely mimicked class A *RHO*-associated human RP clinical hallmarks, which showed early progression of photoreceptor loss. The pathological study of RP in rodents indicates that the accumulation of mutant RHO protein in subcellular compartments may trigger an unfolded protein response (UPR) with cytoprotective outputs, reduce RHO protein synthesis, and result in endoplasmic reticulum (ER)-stress responses, causing rod degeneration[40, 41]. However, the underlying mechanism and pathology of human RP remains unclear. Therefore, our model can offer a useful tool for further mechanistic research on human RP. Furthermore, the characteristics and molecular mechanisms can be extensively studied using our NHP model, facilitating the development of new therapeutic strategies for the disease. In addition, considering that most human RP patients show point mutations, macaque mutants with precise point mutations could be established in the future by base editing, e.g., adenine base editor or cytosine base editor [42], to more accurately mimic human RP disease.

The Cas9-*RHO*-mutant is an acute model for *RHO* pathology based on AAV-mediated gene editing. Considering glial scars were detected in the Cas9-CTRL group, the retinal pathological changes were intermingled with the healing processes from the operation. Nevertheless, the Cas9-*RHO* retinae were generated alongside the Cas9-CTRL retinae in our study. Therefore, the defects in pathological and physiological functions in the Cas9-*RHO* retinae should be due to *RHO* knockout. For future detailed pathogenesis studies, a macaque model with tamoxifen-inducible Cre-ER deletion of the *RHO* mutant could be established. By inducing deletion of *RHO* after complete retinal recovery post injection, the *RHO* deletion-induced phenotypes could be analyzed specifically.

In the Cas9-*RHO*-mutant retinae, progressive photoreceptor degeneration of both rod and cone photoreceptors was detected. As the expression of *RHO* was restricted in rod photoreceptors but not in cone photoreceptors, the loss of cones in Cas9-*RHO* retinae was possibly a secondary effect of the loss of rod photoreceptors, which secrete rod-derived cone viability factor (RdCVF) to protect cones from degeneration [26]. Ciliary neurotrophic factor (CNTF) has also been shown to rescue photoreceptor degeneration in RP models [43, 44]. However, successful examination of CNTF in animal models of neurodegenerative diseases has failed to bring any benefit when tested in human clinical trials. This shows that the efficacy of neurotrophic factors in treating human disease remains to be established in NHP models. As our model recapitulated RP, it provides a reliable tool to test the dosage and safety of neurotropic factors for RP therapy.

The Cas9-*RHO* model also offers unique opportunities to facilitate stem cell transplantation studies. Issues related to immune responses in autologous and allogeneic settings, and the safety or efficacy of either candidate cell lines or cell numbers, can be investigated in NHP models. The NHP model allows assessment of complex physiological, biochemical, behavioral, and imaging end points with direct relevance to human vision. Thus, this study provides a valuable model for human RP studies as well as for the development of novel therapeutic strategies.

Taken together, our study established AAV-delivered CRISPR/SaCas9-mediated gene editing as a rapid and efficient tool for generating NHP disease models, which are advantageous over rodent models. In particular, AAV-delivered CRISPR/SaCas9 can knockout genes in a tissue-specific manner in adult macaques, which has not yet been achieved by germline transmission NHP mutants. This system has broad applicability for editing many other genes or loci to generate human disease models.

## Materials and Methods

### Animals

Four adult male macaques (*Macaca mulatta*), aged 8–13 years, were used in this study based on their similar spectral sensitivities and tri-chromatic color vision as humans. All macaques were maintained at around 25°C on a 12 hour (h) light, 12 h dark schedule. All animal procedures conformed to the requirements of the Institutional Animal Care and Use Committee of the Kunming Institute of Zoology and the Kunming Primate Research Center, Chinese Academy of Sciences (AAALAC accredited). Anesthesia was induced with ketamine (IM, 0.3 g/2 ml, 10–20 mg/kg) and maintained with pentobarbital sodium (IM, 40 mg/ml, 20 mg/kg) for clinical examinations. The pupils of macaques were dilated with tropicamide phenylephrine (Mydrin-P, Santen, Japan) before all examinations.

### Subretinal injection of AAV

Briefly, the adult mice were anaesthetized by intraperitoneal injection with 20 ml/kg 2.5% Avertin (a mixture of 25 mg/ml of Tribromoethylethanol and 25 ml/ml T-amyl-alcohol). After puncture near the limbus, 1.5 μl AAV was injected into subretinal space with a 34G blunted Hamilton needle, passing through the vitreous body and retina. After 3 weeks, the mice were sacrificed and the retinae were cryo-sectioned for fluorescence observation. Adult macaques were anaesthetized with an intramuscular injection of ketamine (10 mg/kg) and pentobarbital sodium (40 mg/kg). Sustained anesthesia was carried out by intravenous dropping with propofol (10–12 mg•kg^-1^•h^-1^) during surgery. Double holes were made near the cornea for entrance of a modified Hamilton needle (38G, 40G) and lighting fiber. The eyes of macaque underwent vitrectomy to lower the intraocular pressure before the addition of extra AAV. Three injection sites were chosen near the macula, superior temporal venule, and inferior temporal venule of the retina (Fig. 3A, right). For each injection site, a retinal swelling was made by the injection of 50 μl of AAV into the subretinal space, passing through the vitreous body and retina.

### Vector construction and AAV production

Annealed guide sequence oligos were inserted to the Bsa1 digested site of pX602 (Addgene plasmid #61593) to construct SaCas9-sgRNA1, sgRNA2, and sgRNA3, respectively. GFP-IRES-PuroR was cloned into the AAV vector (Addgene plasmid #60229) to replace the Cre site for constructing AAV-CBh-GFP, with the same done for AAV-CBh-mKate2. The TBG promoter of pX602 was replaced by human synapsin I promoter. The capsid vector ShH10 was a gift from David V. Schaffer (UC Berkeley). Individual AAVs were produced by transfecting HEK293T cells with three plasmids, i.e., AAV vector, ShH10, and pHelper. The viruses in the nucleus were harvested and purified by iodixanol gradient ultracentrifugation (32 000 rpm, 4 h, 18 °C) in a SW32 rotor, and then concentrated with an Amicon Ultra-15 filter (Millipore). The final titers of AAVs were 5–12 × 10^12^ virus genome/ml, as quantified by qPCR, and all viruses were adjusted to the same titer (5× 10^12^ virus genome/ml) before injection.

### Cell transfection

HEK293T cells were cultured in DMEM supplemented with 10% fetal calf serum (FCS) and 1% penicillin-streptomycin-glutamine (Thermo). Before transfection, the culture medium was replaced with 4% FCS DMEM once the HEK293T cells reached 80% confluence, and polyethylenimine (PEI, Polysciences) was used for cell transfection at a ratio of 2:1 (PEI: plasmid). 6 μg of hSyn-SaCas9/ sgRNA3 plasmid and 6 μg of AAV-CBh-GFP vector were co-transfected for each 6-cm dish. The SaCas9 with scramble sgRNA was used as a control. Puromycin (Thermo) screening (5-8ug/ml) was used to kill untransfected cells, and the living cells were harvested and extracted for genomic DNA (Qiagen kit). Primer-F TTCTCCAATGCGACGGGTGTG and Primer-R AGATAGATGCGGGCTTCCAAC were used to amplify the first exon of *RHO. Apa*I (Thermo) then digested the PCR products for 6 h at 37 °C. The 672-bp band was then cloned into the pEASY-T1 vector (Transgene) for further sequencing.

### PCR and *de novo* sequencing

The *RHO* PCR products were amplified from the genomic DNA (Qiagen kit) of Cas9-CTRL or Cas9-*RHO* infected retinae using Q5 High-Fidelity DNA Polymerase (New England Biolabs) with primer-F1 ACCTTGGGACAGACAAGCCA and primer-R1 TTTCCGAGGGAAACAGAGGC. A shorter and purer 625-bp amplicon was amplified from the second round of PCR using primer-F2 CTTAGGAGGGGGAGGTCACTT and primer-R2 TAACGCTGACAGGAGAGGAGAA. The purified PCR products were *de novo* sequenced and made for a ≤800-bp PCR-free library. The raw data were analyzed by the Burrows-Wheeler-Alignment (BWA) tool to label indel information in the *RHO* PCR products. The indel read numbers for each base pair were counted using a self-written code based on the CIGAR data. The house-developed code was deposited in Github website (https://github.com/Li-Shouzhen/code).

### RT-PCR of SaCas9

The macaque retinae were cut into pieces and lysed by Trizol (Life Technologies, USA), including Cas9-*RHO*- and Cas9-CTRL-infected and uninfected retinae. RNA was extracted and cDNA was synthesized by PCR (PrimeScript RT reagent kit with gDNA Eraser, TaKaRa). SaCas9 was amplified by PCR with Q5 High-Fidelity DNA Polymerase (New England Biolabs) using primers CATCGACTACGAGACACGGG and TTGTGCACGCCTCTTCTCTT. The PCR products were electrophoresed and analyzed.

### Immunohistochemistry and imaging

The mouse and macaque retinae were fixed in 4% paraformaldehyde in phosphate-buffered saline (PBS) (w/v) for 2 h at room temperature and dehydrated by 30% sucrose in PBS (w/v) overnight at 4 °C. Samples were cryosectioned at 14 μm. The slides were rehydrated with PBS and subsequently incubated in blocking buffer (2% BSA, 5% goat serum, 0.5% Triton X-100 in PBS) for 2 h at room temperature. Primary antibodies were incubated with sections overnight at 4 °C. Primary antibodies included rhodopsin (1:1 000, Sigma), sw-opsin (1:500, Millipore), mw/lw-opsin (1:500, Millipore), and GFAP (1:500, Sangon). Slices were washed with PBS three times and incubated with secondary antibodies (1:500, Thermo Scientific) for 2 h at room temperature. DAPI (Thermo Scientific) was used to stain cell nuclei. Confocal images were captured using the Leica SP8 microscope.

### Transmission electron microscopy

The retinae were fixed in 2.5% glutaraldehyde and 2% paraformaldehyde in PBS immediately for 2 h at room temperature. The infected retinae with retinal pigment epithelium (RPE), about 1 mm^3^, were cut and fixed in 2.5% glutaraldehyde and 2% paraformaldehyde overnight at 4 °C. After washing in cacodylate buffer three times, 1.5% (w/v) potassium ferrocyanide, 1% (w/v) osmium tetroxide, or 1% (w/v) osmium tetroxide were individually added for post-fixation twice. 2% (w/v) uranyl acetate in double-distilled water was used for one hour as the first staining before rinsing and dehydrating in ethanol and acetone. The samples were then infiltrated with gradient Epon, and the blocks were hardened for 48 h at 60 °C. Sections (70 nm) were cut with an ultramicrotome (Leica) and collected on formvar/carbon-coated copper grids. The grids were rinsed and dropped in 2% aqueous uranyl acetate, followed by lead citrate for the second staining. Sample grids were photographed using a Hitachi HT7700 transmission electron microscope at 80 kV accelerating voltage.

### Western blot analysis

Samples were lysed by 100 μl of RIPA lysis buffer (150 mM sodium chloride, 1.0% NP-40, 0.5% sodium deoxycholate, 0.1% sodium dodecyl sulfate (SDS), 50 mM Tris, pH 8.0) supplemented with cOmplete and PhosSTOP (Roche), and incubated on ice for 30 min. After 20 min of centrifugation at 12 000 rpm and 4 °C, the supernatant was collected and mixed with 100 μl of loading buffer to produce the final protein sample. The protein extracts were run on a 5%–12% gel and transferred to nitrocellulose (NC) membranes (Millipore). After blocking in a milk solution (5% powdered milk in PBS) for 2 h at room temperature, the membranes were incubated with anti-RHO or anti-ACTIN primary antibodies (Sigma) overnight at 4 °C. The membranes were then washed three times in TBST (0.1M Tris-HCl pH7.5, 0.15 M NaCl, 0.1% Tween-20), and horseradish peroxidase (HRP)-conjugated antibody (1:4 000) in TBST was added for 2 h at room temperature. After washing three times, the membranes were visualized with chemiluminescent substrate (Thermo) on an ImageQuant LAS4000 digital imaging system.

### Clinical examination

An optical coherence tomographic (OCT) device (Spectralis; Heidelberg Engineering, Heidelberg, Germany) was used to image all the retinae before and after surgical injection at different time intervals. Furthermore, fundus fluorescein angiography (FFA) imaging of all animals was carried out by a confocal scanning laser ophthalmoscope (HRA2, Heidelberg Engineering, Heidelberg, Germany) after sodium fluorescein injection under 488-nm laser light excitation. A band-pass filter with a cut-off wavelength of 500 nm was placed in front of the detector. The fundi of macaques was imaged with a fundus camera (Topcon Corp., Japan) before and after surgical injection.

### *Ex vivo* electrophysiology (ERG)

The macaques were dark-adapted for 1 h before *ex vivo* ERG analysis. The eyes were enucleated in dim red light. The retinae with RPE were removed from the eyecup and cut into several pieces, which were then incubated in Ames’ medium (120 mM NaCl, 22.6 mM Na_2_CO_3_, 3.1 mM KCl, 0.5 mM KH_2_PO_4_, 1.5 mM CaCl_2_, 1.2 mM Mg_2_SO_4_, 6 mM glucose, sodium pyruvate, and vitamins) on ice bubbled with 95% O_2_/5% CO_2_ for 30 min dark adaptation before ERG recording. For *ex vivo* ERG recording, retina samples without RPE were transferred to a recording chamber filled with CO_2_ equilibrated Ames’ medium at room temperature (photoreceptor side up). ERG recordings were conducted between a ground electrode in the bottom chamber and a recording electrode positioned above the retina, as shown in Fig. 5A. Flashes (20 ms, 535 nm) were used to stimulate photoreceptors at stepped light intensities from 3.8 to 1 515.6 photons•μm^-2^. The ERG signals were amplified, low-pass filtered at 20 Hz, and digitized for further analysis. After ERG recordings, fluorescent imaging was performed to identify whether the recording region was infected by AAV. For ERG recording in mice, Ames’ solution was supplemented with synaptic transmission blockers (STBs) [20 μM l-(+)-2-amino-4-phosphonobutyric acid (l-AP4) to the solution to block on-bipolar cell signals, 20 μM 6-cyano-7-nitroquinoxaline-2,3-dione (CNQX) to block AMPA/kainate receptors, and 20 μM d-2-amino-5-phosphonovalerate (d-AP5) to block N-methyl-d-aspartate receptors] to exclude b-wave from a-wave.

### Statistical analysis

Quantitative comparisons between experimental groups were analyzed by two-tailed Student’s *t*-test or one-way analysis of variance (ANOVA). Data were presented as mean ± s.e.m. In all cases, *P*-values less than 0.05 were regarded as statistically significant.

## Acknowledgments

We thank Dr. David V. Schaffer from UC Berkeley for the capsid vector ShH10.

## Author Contributions

T.X., X.H., and M.Z. conceived the project and designed the experiments. T.X., X.H., and M.Z. wrote the manuscript with input from all authors. S.L. constructed all plasmids, produced the virus, and finished the phenotypic analysis of models by immunostaining, ERG, and TEM. Y.H. and Y.L. were involved in retinal injection, OCT, and FFA analysis. M.H., W.W., Y.M., Y.C., M.W., Y.Y., Y.W., K.D., Y.G., Z.Q., H.Z., and J.B. were all involved in data analysis and paper discussion.

## Data and material availability

All data needed to evaluate the conclusions in this paper can be found in the manuscript and/or Supplementary Material. Additional data are available from the lead contact (Tian Xue) upon reasonable request.

## Supplementary Figures and Tables

**S1:**
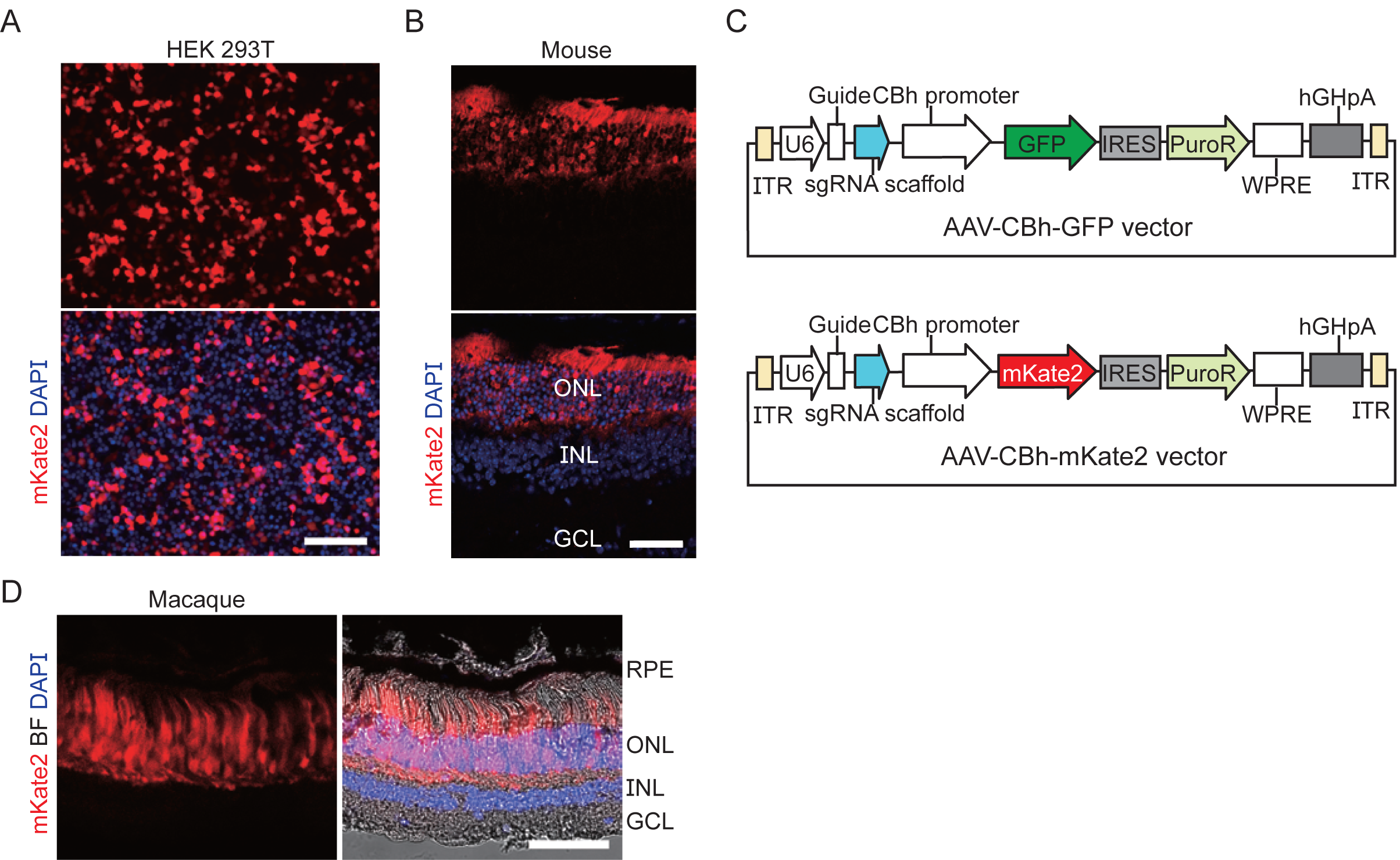
AAV transduction in mouse and macaque retinae. (A) The hSyn promoter drives the expression of mKate2 in HEK 293T cells. Scale bar, 80 μm. (B) Photoreceptors were specifically infected by AAV/ShH10-hSyn-mKate2 in mouse retinae using subretinal injection. Scale bar, 50 μm. (C) Schematic representation of AAV-CBh-GFP and AAV-CBh-mKate2 vectors. (D) Photoreceptors were specifically infected by AAV/ShH10-hSyn-mKate2 in macaque retinae using subretinal injection. Scale bar, 50 μm.

**S2:**
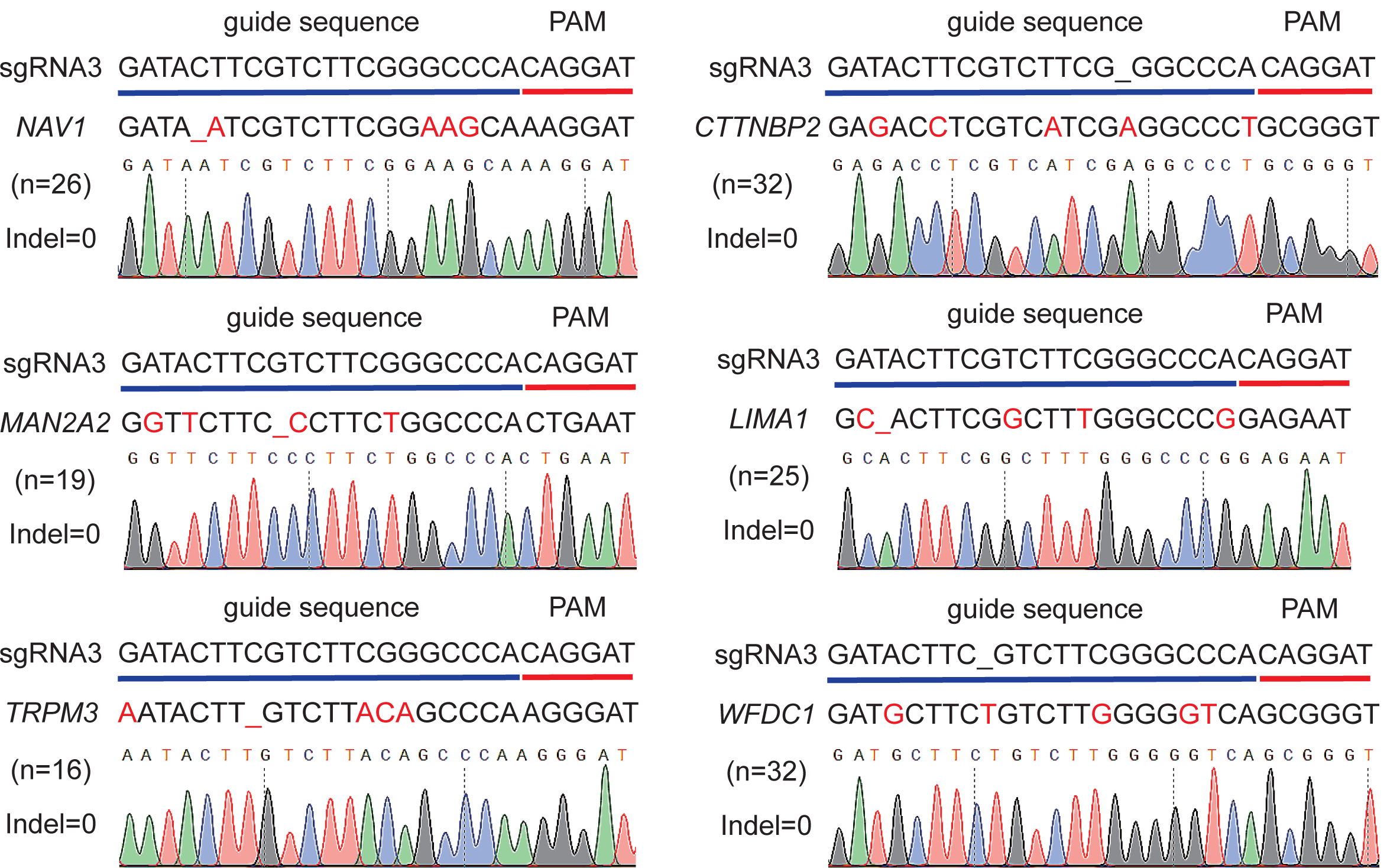
Off-target analysis by PCR product sequencing. PCR product sequencing results at predicted off-target sites. Several sequencing results were shown and compared with wild-type gene sequences.

**S3:**
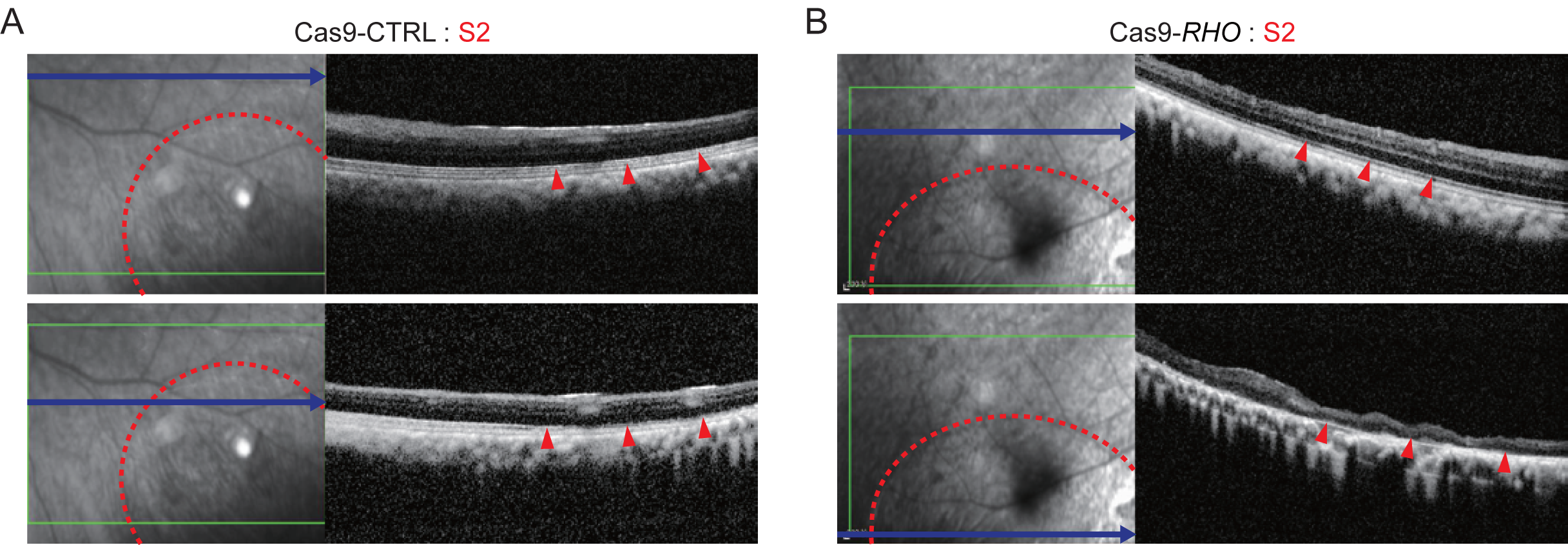
OCT images of S2 injection sites in Cas9-CTRL and Cas9-*RHO* retinae. (A) OCT images of S2 in Cas9-CTRL retinae. Red arrow indicates photoreceptor segments (ellipsoid zone or interdigitation zone). (B) OCT images of S2 in Cas9-*RHO* retinae. Red arrows indicate severe degeneration of outer nuclear layer.

